# Effects of carbon, nitrogen, phosphorus, and potassium on flowering and Fruiting of *Glycyrrhiza uralensis*

**DOI:** 10.1101/2021.01.23.427937

**Authors:** Binbin Yan, Yan Zhang, Xiaobo Zhang, Sheng Wang, Jie Cui, Kai Sun, Tielin Wang, Chuanzhi Kang, Jiahui Sun, Yang Ge, Lanping Guo, Wenquan Wang

## Abstract

**Abstract:** Carbon (C), nitrogen (N), phosphorus (P), and potassium (K) play an important role in flower bud differentiation and seed-filling; however, the effects of these elements on the flowering and fruiting of *Glycyrrhiza uralensis* Fisch. are not known. In this study, we evaluated the differences in the C, N, P, and K levels between the fruiting and nonfruiting plants of *G. uralensis* at different growth stages. The correlations between the elements C, N, P, and K and the flower and fruit falling rates, rate of empty seeds, rate of shrunken grains, and thousand kernel weight (TKW) were also determined. The results show that the P and K levels and C:N, P:N, and K:N ratios of flowering plants are significantly higher than those of nonflowering plants; N level of flowering plants is significantly lower than that of nonflowering plants at the flower bud differentiation stage. The number of inflorescences was positively correlated with C and K levels and C:N and K:N ratios. A low level of C, P, and K and high level of N in flowering and pod setting stage may lead to the flower and fruit drop of *G. uralensis*. The K level is significantly negatively correlated with the rates of empty and shrunken seeds. The N level is significantly positively correlated with TKW. Thus, high levels of C, P, and K might be beneficial to flower bud differentiation, while higher levels of N is not beneficial to the flower bud formation of *G. uralensis*. Higher levels of N and K at the filling stage were beneficial to the seed setting and seed-filling of *G. uralensis*.

**Highlight:** High levels of C, P, and K might be beneficial to flower bud differentiation, while higher levels of N is not beneficial to the flower bud formation of *G. uralensis*. Higher levels of N and K at the filling stage were beneficial to the seed setting and seed-filling of *G. uralensis*.

## Introduction

*Glycyrrhiza uralensis* Fisch., a perennial herb of the Leguminosae family (Chinese Pharmacopoeia Commission, 2020), is a common bulk herb in China, and it has been widely used in the medical, food, tobacco, fodder, and cosmetic industries (Seki et al., 2011; Vaya et al., 1997; Wang et al., 2011). Wild *G. uralensis* has been the main source of licorice for decades. Overharvesting has gradually exhausted wild *G. uralensis* resources, and therefore cultivated *G. uralensis* has become an alternative source. Although *G. uralensis* exhibits seed propagation, its seed yield is low, and its fruiting rate under natural conditions is only 10–21% (Wang et al., 2003). Thus, the current *G. uralensis* production fails to satisfy the industrial demand. Flower bud differentiation and seed-filling are essential in the fruiting of plants. The elements C, N, P, and K play an important role in these processes. However, the effects of C, N, P, and K on the flowering and fruiting of *G. uralensis* are unknown.

At the beginning of the 20th century, Klebs systematically studied the changes in various components during plant flowering. The proportion of carbohydrates (C) and available nitrogen compounds (N) played a key role in the flowering of plants. Therefore, Klebs proposed the famous theory carbon nitrogen ratio (C:N). Higher the C:N ratio, higher the plant reproductive growth; otherwise, higher the plant vegetative growth(F, 2000; W, 1967). However, a large number of subsequent studies have shown that the C:N ratio theory cannot fundamentally explain the essence of plant flower formation, but it has a significant effect in controlling flower bud differentiation. As a key regulator of plant flowering, N plays an important regulatory role in plant flowering(Miyazaki et al., 2014): N can promote the differentiation of floret primordia(Miyazaki et al., 2014). C is an important material basis of plant life metabolism, promoting floret development(Mi et al., 2005). A higher C:N ratio was found to be beneficial to the flower bud differentiation of *Myrica rubra* and radish(Sun et al., 2010; Xu et al., 2009), and the starch and soluble sugar content in potato leaves had an important effect on flowering(Ai et al., 2017). In addition to C and N, phosphorous (P) and potassium (K) also play an important role in plant flowering and fruiting. P is one of the essential nutrients for plant growth and development(Zhang et al., 2013). An appropriate P content could regulate the growth and flowering of *Chrysanthemum morifolium*(Wang et al., 2012), and an appropriate K content could significantly promote the flowering of rice(Wei et al., 2019). In addition, nutritional imbalance is also an important cause of crop flower and fruit drop(Zhang et al., 2016).

The elements C, N, P, and K also significantly affect seed development. A low content of C was is the cause of insufficient grain filling in rice(You et al., 2017), and a higher C content in coffee leaves is conducive to its flowering(Lin et al., 2019). A lower N content is not conducive to the grain development of millet(Cheng et al., 2016). K plays an important role in wheat grain filling, increasing the supply of soluble sugars and sucrose in seeds and accelerating the accumulation of starch(Wang et al., 2003). P can promote the accumulation of reducing sugar, sucrose, soluble sugar, and starch in cucumber seeds(Zhang et al., 2018).

Higher P:N and K:N ratios are beneficial to ginkgo fruit development(Sha, 2006)。

This study aims to explore the relationship between C, N, P, and K levels and ratios and the flowering and fruiting of *G. uralensis* by comparing the differences in C, N, P, and K content in flowering fruiting and nonflowering licorice plants at different growth stages and ages. Furthermore, the dynamic changes in the C, N, P, and K levels at different growth stages, correlations between the C, N, P, and K levels and ratios with the number of inflorescences, flower and fruit falling rates, and fruiting rate were analyzed. This study not only provides a theoretical reference for interpreting the flowering mechanism of *G. uralensis*, but also lays a theoretical foundation for the control of seed yields.

## Materials and methods

### Experimental Materials

*G. uralensis* plants in different growth years were used. The test site is in Hedong Town, Guazhou County, Jiuquan City, Gansu Province (40°31′N, 96°42′E), where the altitude is approximately 1378 m. The mean annual precipitation is 83.5 mm. The maximum, minimum, and annual average temperatures are 36.7 °C, −17.6 °C, and 8.75 °C, respectively. The annual sunshine duration is 3237 h.

### Experimental Design

Two-, three-, five-, seven-, and 12-year-old *G. uralensis* plants were selected. Three sampling plots (5 × 5 m^2^) were randomly selected in the test field for three repetitions. The study involved the following four experimental designs:

Two-, three-, and seven-year-old *G. uralensis* plants were considered as the subjects. In September of the first year, 100 plants were stake-marked in quadrats of various ages. The buds of the plant labeled were taken on 1 May of the following year, and the fifth compound leaves at the top of each labeled plant on 21 May, 17 July, and 13 September were selected and stored in the corresponding numbered envelope, killed green at 105 ☐, and then dried at 80 ☐.

On May 10, 100 plants numbered 1–100 in the quadrat were listed and labeled as five-, seven-, and 12-year-old samples. The fifth compound leaf at the top of each marked plant was obtained from the budding stage (May 11), squaring stage (May 21), seed-filling stage (July 17), and seed maturation stage (September 13), killed green at 105 ☐, and then dried at 80 ☐. On September 13, the collected samples were grouped and merged according to the flowering and fruiting conditions. Each growth year was divided into a flowering and fruiting group and a nonflowering group.

The sampling time for each period was between 8 AM and 9 AM.

On June 18, (full bloom), 100 plants were randomly labeled in the five-, seven-, and 12-year-old quadrats, the inflorescence number of each plant was counted, and the flower and fruit abortion rates of each plant were counted on 17 July (the filling stage). The empty seed rate and shrinkage rate of each plant were counted after 13 September, and the thousand kernel weight (TKW) of seeds in each sampling plot was tested at the same time.

### Determination of C, N, P, and K Content

First, the sample was initially crushed, and then two sufficient samples of coarse powder were selected using quartering method. One part of the powder was further crushed to pass through a 100 mesh sieve, and a sufficient amount of powder was selected using quartering method to determine the total C content in the sample. The other part was continued to be crushed and passed through a 60 mesh sieve. A sufficient amount of sample was selected using quartering method to determine the N, P, and K contents in the sample.

The total C content was determined using oil bath method. The total N content was determined using Kjeldahl method in a mixed CuSO_4_-K_2_SO_4_ digestion solution. The sample was digested using H_2_SO_4_-H_2_O_2_ method; the decocting solution was used to determine the content of P and K; the total P content was determined using a ultraviolet spectrophotometer; the total K extract was determined using an AA flame photometer.

### Statistical Analysis

SPSS 19.0 was used for data descriptive statistics, factor correlation analysis, and variance analysis (i.e., ANOVA). The contents of endogenous hormones in the flowering and nonflowering plants in each sample were determined three times, and variance analysis was performed using the average value of each sample. The data obtained from each sampling date were analyzed separately using the Duncan Analysis Method, and the resulting means were tested using the least significant difference test at the P0.05 level (LSD0.05).

## Results

### Changes in C, N, P, and K levels during growth of *G. uralensis*

By comparing the changes in the C, N, P, and K levels and ratios in two-, three-, and seven-year-old plants at different stages, the changes in C, N, P, and K during plant growth were analyzed. The results (Figure 1) show that the changes in the plant C, N, P, and K levels are the same in different years. At the germination stage (May 1), the C level was low (150–180 g/kg), and then it increased gradually with vigorous life activities. At the seedling stage (May 11), the C level increased rapidly to a higher level (370–390 g/kg). It increased by 51%, 59%, and 67% in two-, three-, and seven-year old plants, respectively. At the vigorous-growth stage of the aboveground part, the C level reached the highest level (444–446 g/kg) and then gradually decreased (Figure 1A). At the germination stage, the N level was low (6.5–8.2 g/kg), rapidly reached to the highest level (16–18 g/kg) at the seedling stage, and then gradually decreased. At the end of growth stage of the aboveground part, the N level decreased rapidly. The N level of aboveground part decreased to 6–8 g/kg in the senescence stage (Figure 1B). The P content of plant remained at a high level from the germination stage to the end growth stage of the aboveground part and gradually decreased after the stop growth of the aboveground part (Figure 1C). At the germination stage, the K level was low (8–10 g/kg) and then gradually increased with vigorous life activities. It rapidly reached to a higher level at the seedling stage and then remained at a higher level (15–16 g/kg) and gradually decreased after the stop growth of aboveground part (Figure 1D).

**Figure 1.**
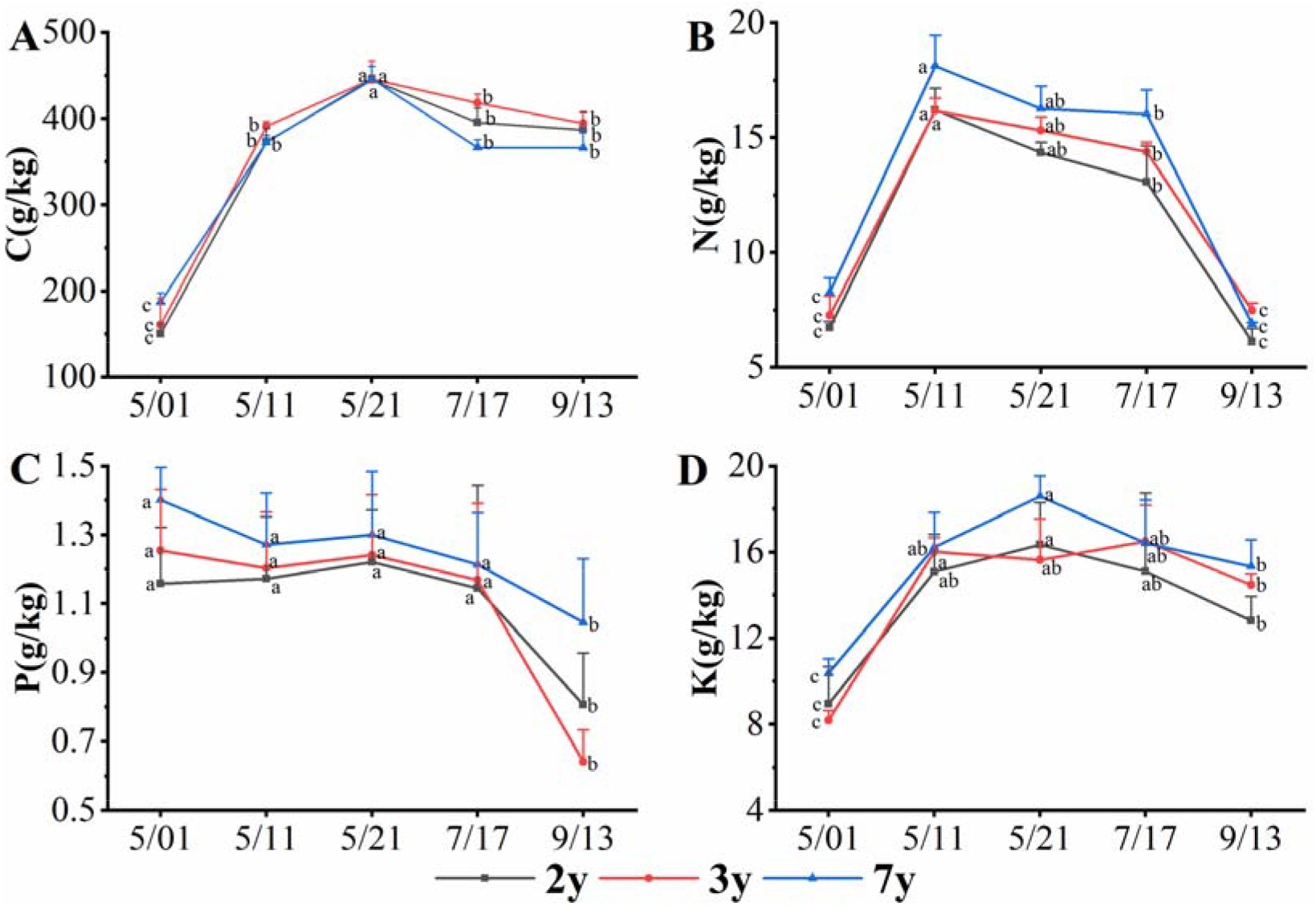
Changes in the C, N, P, and K levels throughout the growth of *G. uralensis*. Different letters reflect significant differences among different indexes in the same stage: (A) C content, (B) N content, (C) P content, and (D) K content.

The C:N ratio rapidly increased from the germination stage to the autumn when growth stopped (Figure 2A). The P:N ratio rapidly decreased from the germination stage to the vigorous-growth stage of aboveground parts and then gradually increased until autumn when growth stopped (Figure 2B). The K:N ratio remained at a low level from the germination stage to the vigorous-growth stage and then rapidly increased until autumn when growth stopped (Figure 2C).The variation in C:P ratio is similar to that of C:N ratio (Figure 2D), but the variation in P:K ratio is opposite (Figure 2E).

**Figure 2.**
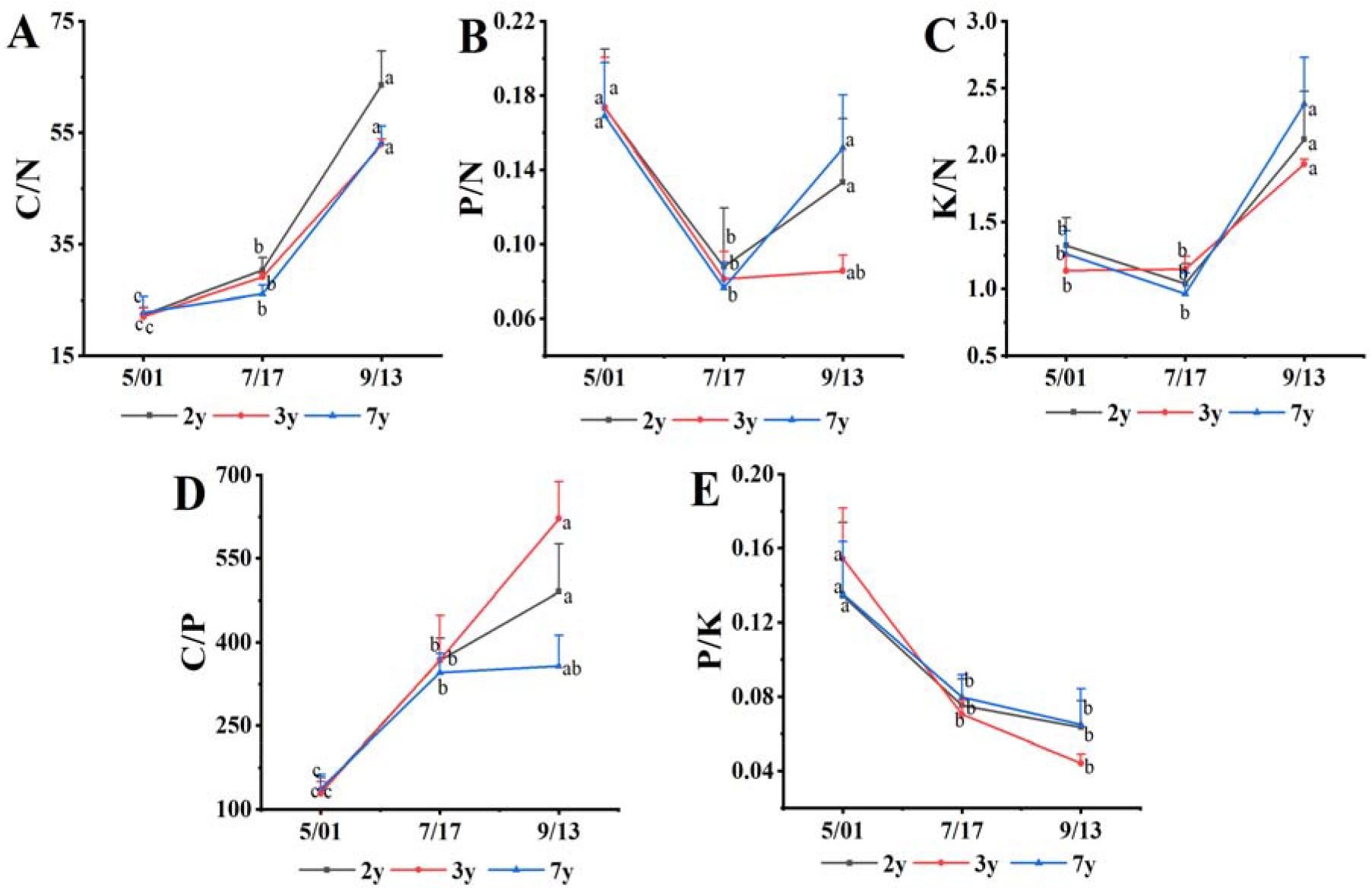
Changes in C, N, P, and K ratios during the growth of *G. uralensis*. The different letters reflect significant differences among different indexes in the same stage e: (A) C:N ratio, (B) P:N ratio, (C) K:N ratio, (D) C:P ratio, and (E) P:K ratio.

The lowest C, N, and K levels were obtained in the germination stage and remained at a high level at the growth stage of aboveground parts. After the aboveground part stopped growing, the C, N, P, and K levels decreased gradually. The C, N, P and K levels decreased gradually when the growth of aboveground part stopped.

### Relationship between C, N, P, and K levels and flower bud differentiation of *G. uralensis*

The effects of C, N, P, and K levels on the flower bud differentiation of licorice were analyzed by comparing the differences in the C, N, P, and K levels among five-, seven-, and 12-year-old flowering and fruiting plants and nonflowering plants during the flower bud incubation (May 11) and bud stage (May 21). The results (Figure 3) show that the difference trends of the C, N, P, and K contents and ratios between the five-, seven-, and 12-year-old flowering and nonflowering plants are the same. The N level of five-, seven-, and 12-year-old nonflowering plants at seedling stage and bud stage is significantly higher than those of flowering plants by 23%, 19%, 24% and 71%, 113%, 73% respectively (Figure 3A). The P level of flowering plants at seedling stage and bud stage is significantly higher than those of nonflowering plants (Figure 3B). No significant difference in K content was observed between the flowering and nonflowering plants at seedling stage, but it was significantly higher than those of nonflowering plants at the flower bud stage by 45%, 17%, and 54% of five-, seven-, and 12-year-old plants, respectively (Figure 3C). No significant difference in C level was observed between the flowering and nonflowering plants at different stages.

**Figure 3.**
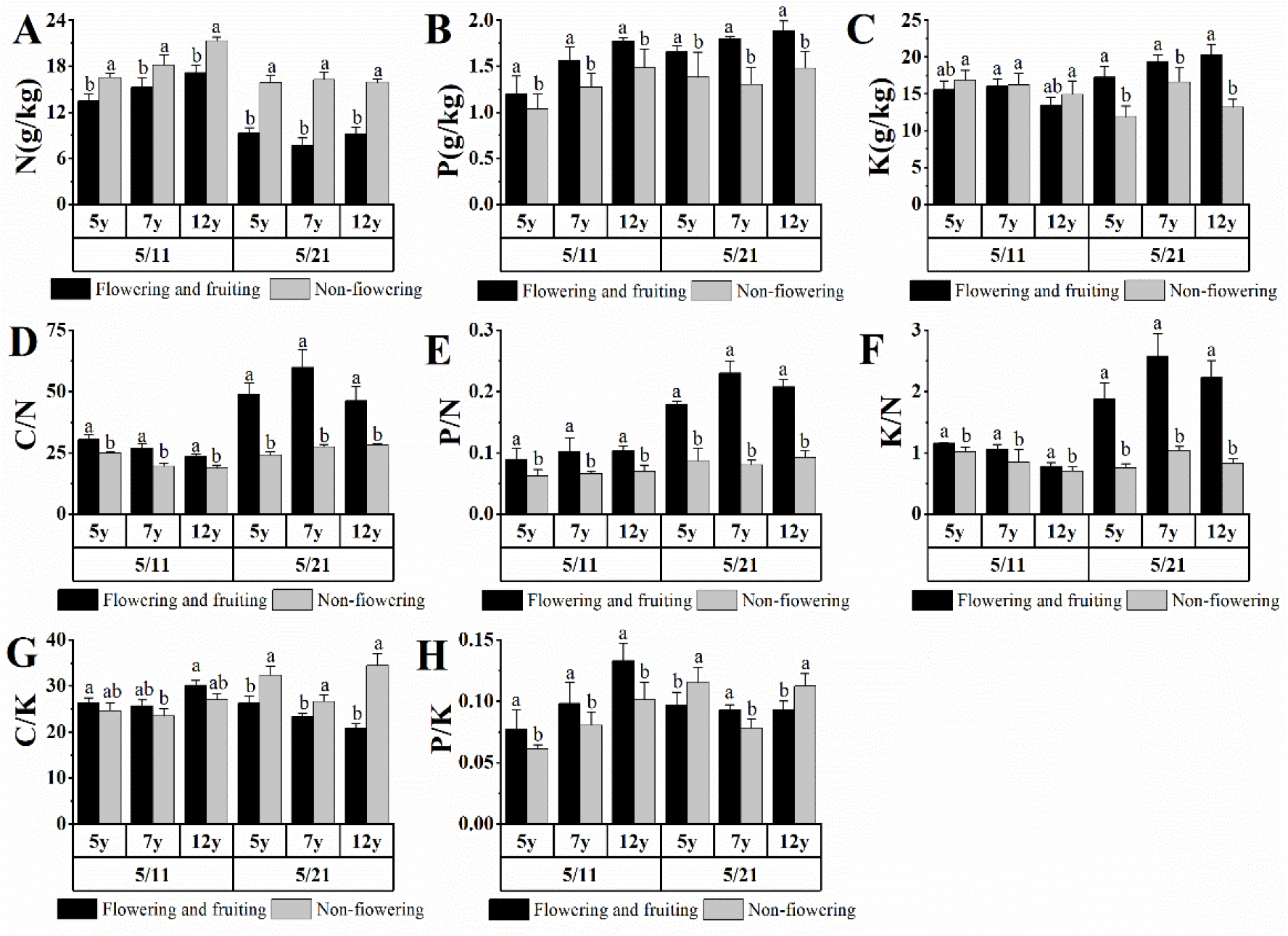
Effects of C, N, P, and K contents and ratios on *G. uralensis*. flower bud formation. Different letters reflect significant differences among different indexes in the same stage: (A) N content, (B) P content, (C) K content, (D) C:N ratio, (E) P:N ratio, (F) K:N ratio, (G) C:K ratio, and (H) P:K ratio.

The C:N, P:N, and K:N ratios of five-, seven-, and 12-year-old flowering plants at seedling stage and bud stage were significantly higher than those of nonflowering plants by 29%, 49%, 17% and 95%, 138%, 155%, respectively (Figures 3D, E, F). The C:K ratios of five-, seven-, and 12-year-old flowering plants at seedling stage were significantly higher than those of nonflowering plants, but it was significantly lower than that of nonflowering plants at flower bud stage (Figure 3G). The P:K ratios of five-, seven-, and 12-year-old flowering plants at seedling stage were significantly higher than those of nonflowering plants (Figure 3H).

All the above data indicated that the elements C, N, P, and K may be related to the flower bud differentiation of *G. uralensis*. The higher levels of P and K levels and C:N, P:N, and K:N ratios and lower levels of N in bud stage may benefit the flower bud differentiation in *G. uralensis*.

The average number of inflorescences per plant is different among different sampling plots (Table S1), ranging between 3.53 and 5.65. The correlation between the number of inflorescence and contents of C, N, P, and K was analyzed. The results are shown in Table 1. The number of inflorescences showed an significantly positive correlation with C contents, as well as C:N and K:N ratios, and extremely significantly with K contents.

**Table 1.**
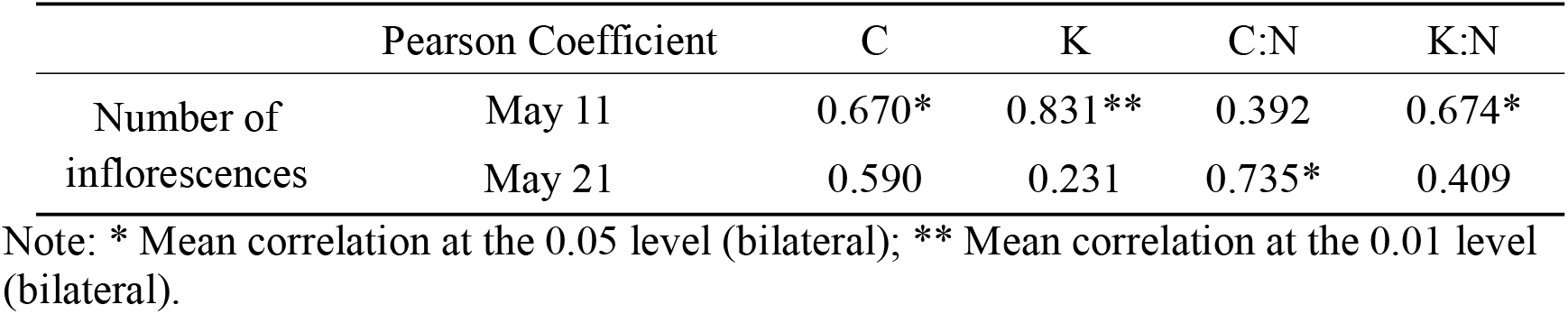
Correlation between C, N, P, and K levels and number of inflorescence (n = 9)

The results showed that high C, P, and K levels benefited the flower bud differentiation of *G. uralensis.*

### Differences in C, N, P, and K levels between fruiting and nonflowering plants at different stages

The 100 selected five-, seven-, and 12-year-old plants covered both fruiting and nonflowering plants (Table S2). In Figure 4, the C, N, P, and K contents of the flowering, fruiting, and nonflowering plants are the average contents of the five-, seven-, and 12-year-old flowering and fruiting plants, as well as nonflowering plants. This study aimed to compare the differences in the C, N, P, and K levels among the flowering and fruiting plants and nonflowering plants at different stages.

**Figure 4.**
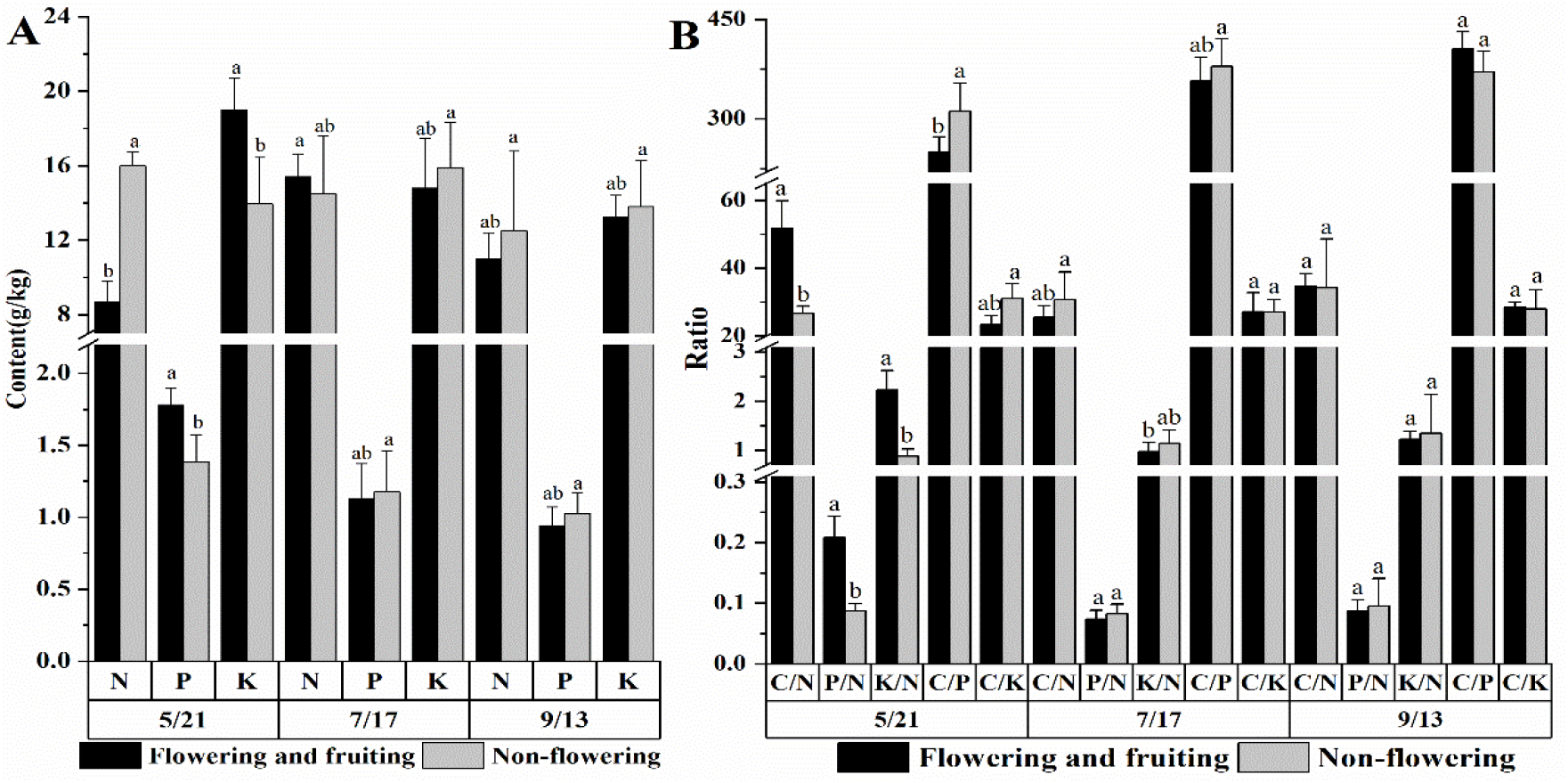
Differences in C, N, P, and K levels between flowering and fruiting plants and nonflowering plants at different growth stages. The different letters reflect significant differences among different indexes in the same stage. (A) N, P, and K content, (B) C:N, P:N, K:N, C:P, and C:K ratios.

The P and K levels of flowering plants at bud stage were significantly higher than those of nonflowering plants by 28% and 36%, respectively, and the N levels were significantly lower than those of nonflowering plants. No significant difference was observed in the C, N, P, and K contents between flowering and nonflowering plants from the seed-filling stage to the seed maturity stage (Figure 4A).

The C:N, P:N, K:N, C:P, and C:K ratios of flowering plants at bud stage were significantly higher than those of nonflowering plants by 93%, 140%, 154%, 24.92%, and 32.64%, respectively, and the N levels were significantly lower than those of nonflowering plants. No significant difference was observed in the C, N, P, and K contents between flowering and nonflowering plants from the seed-filling stage to seed maturity stage (Figure 4B).

All these data indicate that the C, N, P, and K levels may be related to the flower bud differentiation of *G. uralensis*. The higher P and K levels and C:N, P:N, and K:N ratios and lower N levels in bud stage may benefit the flower bud differentiation in *G. uralensis*.

All the above data indicate that the C, N, P, and K levels may be related to the flowering and fruiting of *G. uralensis*. The high P and K levels and C:N, P:N, K:N, C:P, and C:K ratios may be beneficial to the flowering and fruiting of licorice, and a high N level may be unfavorable to the fruiting of *G. uralensis*.

### Correlation between C, N, P, and K levels and flower and fruit falling rates of *G. uralensis*

Flower and fruit falling covers two situations, namely inflorescence falling, which was observed in some plants in all sampling plots (Table S2), and floret or pod falling (Figure S1A; Table S3). To establish the relationships between the C, N, P, and K contents and flower and fruit falling, we analyzed the differences in C, N, P, and K contents between the fruiting plants and inflorescence falling plants from the flower bud stage to the seed-filling stage. Correlations between the flower and fruit falling rates and C, N, P, and K contents were analyzed.

The results (Figure 5) show that the N level of exfoliated inflorescence plants was considerably higher than those of flowering and fruiting plants at flower bud stage (*p* < 0.05); No significant difference was observed in the C, N, P, and K contents between inflorescence falling plants and flowering and fruiting plants. The P, K levels and C:N, P:N, K:N, C:P, and C:K ratios of flowering and fruiting plants at bud stage were significantly higher than those of inflorescence falling plants (*p* < 0.05), and no significant difference was observed at the seed-filling stage. The rates of falling flowers and fruit were negatively correlated with the C and P levels at the bud stage, and there was no significant correlation with the C, N, P, and K contents in other stage (Table 2).

**Figure 5.**
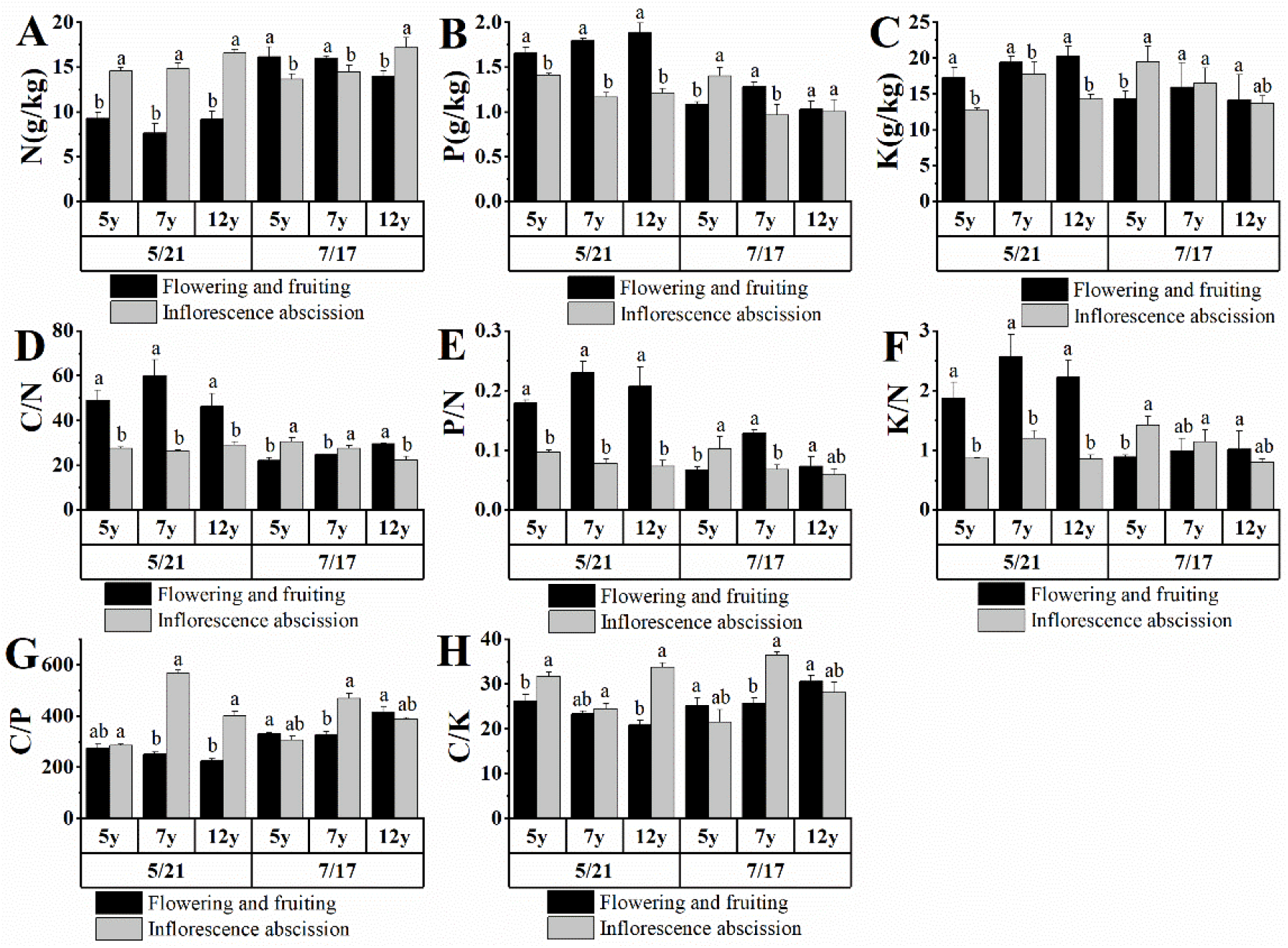
Correlation between C, N, P, and K levels and flower and fruit falling rates of *G. uralensis.* The different letters reflect significant differences among different indexes in the same stage: (A) N content, (B) P content, (C) K content, (D) C:N ratio, (E) P:N ratio, (F) K:N ratio, (G) C:P ratio, and (H) C:K ratio

**Table 2.** Correlation between C, N, P, K and the rate of falling flowers and fruits of *G. uralensis* (*n* = 9)

| Pearson Coefficient | Date | C | P |
| --- | --- | --- | --- |
| Rates of falling flowers and fruit | May 21 | -0.842** | -0.711* |

Note: \* Mean correlation at the 0.05 level (bilateral); \*\* Mean correlation at the 0.01 level (bilateral).

**Table 3.**
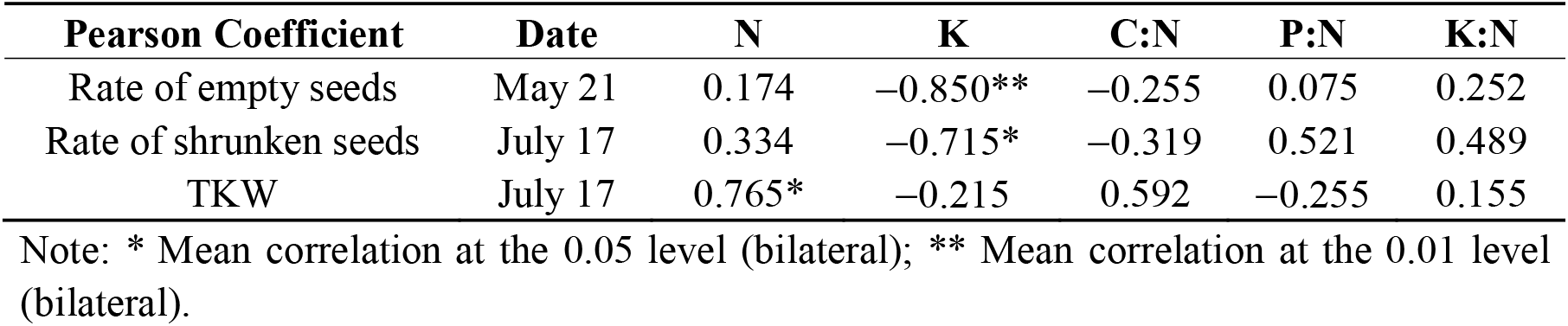
Correlation of C, N, P, and K levels with rate of empty seeds, rate of shrunken seeds, and TKW of *G. uralensis* (*n* = 9)

These results show a strong correlation between the C, N, P, and K contents and the flower and fruit falling of *G. uralensis*. The higher N level and lower P and K levels and C:N, P:N, K:N, C:P, and C:K ratios are possible causes of licorice inflorescence shedding. Low C and P levels may be the cause of licorice flower and fruit loss.

### Correlations of C, N, P, and K levels with the number of empty seeds, number of shrunken seeds, and TKW

The correlations between the C, N, P, and K contents and ratio with the rate of empty seeds, rate of shrunken seeds, and TKW in different sampling plots (Table S4; Figure S1B,C) were analyzed (Table 2).

The rate of empty seeds was extremely significantly negatively correlated with the K content on May 21 (bud stage) (*p* < 0.01). The rate of shrunken seeds showed a significantly positive correlation with the K content on July 17 (seed-filling stage). During the same period, TKW was significantly positively correlated with the N content (*p* < 0.05).

These data show a strong correlation between the N and K content and the fructification of *G. uralensis.* A high level of K content in flower bud stage, seed-filling stage, and mature stage and a high level of N content from seed-filling stage to mature stage are beneficial to the seed setting of *G. uralensis*.

## Discussion

The C:N ratio of flowering and fruiting plants was significantly higher than that of nonflowering plants (*P* < 0.05). A higher C:N ratio might be beneficial to the flower bud differentiation of *G. uralensis*. The results are similar to those of black grape. A decrease in the content of total N, increase in the content of total C, and increase in the C:N ratio would shorten the time of flower bud differentiation of black grape(Zhao et al., 2016), and the increase in C:N ratio is conducive to the flower bud differentiation of Chinese prickly pepper and summer maize(He et al., 2018; Li et al., 2012). The higher C:N ratio was beneficial to wheat seed setting, in which N promoted the differentiation of floret primordium, and C promoted floret development(Ni et al., 2013). The N content of five-, seven-, and 12-year-old flowering plants was significantly lower than that of nonflowering plants at both bud formation and budding stages, but no significant difference was observed in the C content. Therefore, the effect of N on the flowering of *G. uralensis* is more obvious than that of C. The N content in the leaves of flowering branches of litchi was significantly lower than that of nonflowering branches(Zhang et al., 2016), and the N content in the branches and stems of nonflowering *Dendrocalamus latiflorus* was significantly higher than that of flowering plants(Xu et al., 2017). The N utilization rate of cherry was the highest at the flower bud differentiation stage(Zhao et al., 2008).

N accumulation in legumes is one of the main determinants of crop yield. The TKW of *G. uralensis* was significantly positively correlated with the N level at the filling stage and mature stage. It was concluded that the higher level of N content in the filling stage and mature stage was beneficial to the seed filling of *G. uralensis*. An increase in N content improved the seed quality of *Anemone nemorosa* and wheat(Bhatta M, 2017; De Frenne et al., 2018). Faba bean began to accumulate N at a high rate only at the late vegetative growth stage and continued to enter the seed filling at a high rate(Marrou H, 2017), and the autophagy gene *GmATG8c* may be related to this process(Islam et al., 2016). Maize ear was shorter and biomass accumulation was less under N deficiency, and this negative change can be attributed to the significant decrease in nitrate reductase, glutamine synthetase, sucrose phosphate synthetase, and sucrose synthetase activities in N-deficient maize ears(Yu J, 2016). Because cell division occurs between the flowering and beginning of seed filling, the accumulation of C and N reserves in seeds before the beginning of seed filling will affect the growth rate of a single seed, thus affecting the weight of a single seed(Munier-Jolain N, 2008). The seeds of legumes are rich in protein and have a high demand for N. The reactivation of N in the leaves will reduce the photosynthetic activity and shorten the duration of filling period, thus decreasing the weight of individual seeds(Dumas and Rogowsky, 2008).

Based on the results of this study, an appropriate N content is conducive to the differentiation of licorice flower buds and seed filling. Therefore, it can be inferred that during cultivation, a proper control of application amount of N fertilizer is beneficial to the increase of licorice seed yield. Studies have shown that increased application of N fertilizers in different ecological regions increased the wheat yield(Wu et al., 2017). N treatment enhances the storage protein content, endosperm protein body quantity, and partial processing quality by altering the expression levels of certain genes involved in protein biosynthesis pathways and storage protein expression at the proteomics level(Yu et al., 2017). Application of N fertilizer significantly increased the content of protein N, total amino acids, and partially saturated and unsaturated fatty acids in peony seeds for oil(Jiang et al., 2016). The results of this study show that the N content of flowering and fruiting licorice plants during the flower bud differentiation stage is significantly lower than that of nonflowering plants, indicating that excessive N levels at the flower bud differentiation stage are not conducive to the flowering and fruiting of licorice. Therefore, the application time and amount of N fertilizer should be controlled. Studies have shown that a certain amount of N fertilizer can promote the growth and development of sunflower, but excessive N fertilizer is not conducive to its fruiting(Wang, 2016).

During flowering and fruiting, in addition to N, P and K also play an important role(Aguirre et al., 2018; Mesejo et al., 2019). Higher levels of P and K and C:N, P:N, and K:N ratios were beneficial to the flower bud differentiation of *G. uralensis*., and lower C and P levels may be the cause of flower pod and pod abscission of *G. uralensis*. The K level is significantly negatively correlated with the rates of empty and shrunken seeds. Therefore, in production practice, in addition to controlling the application of N fertilizer, it is also necessary to control the application of phosphate and potash fertilizers.

Studies have shown that an increased application of K fertilizer can significantly increase the yield and quality of apples and sunflowers(Tu et al., 2017; Wang, 2016), and the yield of wheat could be significantly increased by applying P fertilizer(Akhtar et al., 2016). Among them, K fertilizer increased the IAA, GA, and ZR contents and decreased the ABA content(Tu et al., 2017). Our research shows that IAA, GA, and ZR are beneficial to the flower bud differentiation and seed filling, while ABA was not beneficial(Yan et al., 2019). Therefore, N, P, and K fertilizers may affect the seed yield by affecting the content of endogenous hormones in licorice plants. A reasonable NPK application promoted flower bud differentiation by affecting the endogenous hormone levels of *Camellia oleifera*(Luo et al., 2019). In summary, according to the results of this study, in the regulation of licorice flowering and fruiting in the future, attention should be paid to the combined application of N, P, and K. Studies have shown that the application of N, P, and K fertilizers improved the quality of rice, such as increasing the rate of heading rice, reducing chalkiness, rice length-to-width ratio, and amylose content of polished rice(Wang et al., 2011). Formula fertilization requires less N but can achieve a higher yield, and can significantly increase the number of rice ears and number of grains per ear(Zhou et al., 2011). Appropriate N, P, and K fertilizer application can significantly increase the seed yield of *Atractylodes macrocephala* and reduce its empty shrinkage rate(Peng, 2016).

## Conclusions

There is a certain relationship between C, N, P, and K contents and the flowering and fruiting of *G. uralensis*. The P and K levels and C:N, P:N, and K:N ratios of flowering and fruiting plants at the seedling stage and flower bud stage were significantly higher than those of nonflowering plants, and the N level at the bud stage was significantly lower than that of nonflowering plants (*P* < 0.05). This shows that in the seedling stage and flower bud stage, higher levels of P and K and C:N, P:N, and K:N ratios and lower levels of N at the bud stage are beneficial to the differentiation of licorice flower buds. Higher levels of N, lower levels of P and K and C:N, P:N, and K:N ratios may be the cause of shedding of licorice inflorescence; lower levels of C and P may be the reasons for the abscission of flower and pod. The empty seed rate was significantly negatively correlated with the K level in the bud stage (*P* < 0.01), and the deflated seed rate was significantly negatively correlated with the K level in the filling stage (*P* < 0.05). TKW was significantly positively correlated with the N level at the seed-filling stage (*P* < 0.05). This shows that a higher level of K content in the flower bud stage and seed-filling stage and a higher level of N content in the seed-filling stage are beneficial to the seed setting and seed filling of *G. uralensis*. Therefore, a proper control of C, P, K, and N contents can facilitate flower bud differentiation and seed-filling, increase the number of inflorescence and TKW, and decrease the rate of flower and fruit falling, empty seeds, and shrunken seeds. These effects ultimately increase the seed yield of *G. uralensis*.

## Supplementary data

Supplementary materials are available online.

Figure S1. Flower and fruit falling, empty seed plants, and licorice seeds;

Table S1. Number of inflorescences in different sampling plots;

Table S2. Flowering and fruiting situations of plants in different sampling plots;

Table S3. Rate of flower and fruit falling in different sampling plots;

Table S4. Rate of empty seeds, rate of shrunken seeds, and TKW in different sampling plots.

## Acknowledgements

Thanks to farmer Zhou Xuede for selflessly providing experimental materials. Thanks to resource center of traditional chinese medicine, chinese academy of traditional chinese medicine and institute of medicinal plants of chinese academy of medical sciences for providing laboratories and other experimental materials to support the experiment.

## Author contribution

Binbin Yan conceived investigation and writing – Original Draft Preparation. Lanping Guo and Wenquan Wang conceived Conceptualization. Yan Zhang conceived data Curation. Xiaobo Zhang conceived funding acquisition. Sheng Wang and Jie Cui conceived Methodology. Kai Sun and Tielin Wang conceived Supervision. Chuanzhi Kang and Jiahui Sun conceived resources. Binbin Yan and Yang Ge conceived Writing – Review & Editing

## Data availability statement

All data supporting the findings of this study are available within the paper and within its supplementary materials published online. (For data available on request from the authors).

The data supporting the findings of this study are available from the corresponding author, (Binbin Yan), upon request.

## References

Chinese Pharmacopoeia Commission. 2015, Pharmacopoeia of the People’s Republic of China, Volume I; China Medical Science Press: Beijing, China, p. 86–87. (In Chinese)

Aguirre, M., E. Kiegle, G. Leo, and I. Ezquer. 2018, Carbohydrate reserves and seed development: an overview: Plant Reprod, v. 31, p. 263–290.

Akhtar, M., M. Yaqub, A. Naeem, M. Ashraf, and V. E. Hernandez. 2016, Improving phosphorus uptake and wheat productivity by phosphoric acid application in alkaline calcareous soils: J Sci Food Agric, v. 96, p. 3701–7.

Bhatta M, R. T. R. D. 2017, Genotype, environment, seeding rate, and top◻dressed nitrogen effects on end◻use quality of modern Nebraska winter wheat: J, Food Agric, v. 97, p. 5311–5318.

De Frenne, P., H. Blondeel, J. Brunet, M. M. Caron, O. Chabrerie, M. Cougnon, S. Cousins, G. Decocq, M. Diekmann, B. J. Graae, M. E. Hanley, T. Heinken, M. Hermy, A. Kolb, J. Lenoir, J. Liira, A. Orczewska, A. Shevtsova, T. Vanneste, and K. Verheyen. 2018, Atmospheric nitrogen deposition on petals enhances seed quality of the forest herb Anemone nemorosa: Plant Biol (Stuttg), v. 20, p. 619–626.

Dumas, C., and P. Rogowsky. 2008, Fertilization and early seed formationFécondation et développement précoce de la graine: Comptes Rendus Biologies, v. 331, p. 715–725.

F, Z. J. W. 2000, The Effect of Supplying Nitrate at Different Seasons on the Growth, Blossoming and Nitrogen Content of Young Apple Tree in Sand Culture: Journal of Jilin Agricultural University.

Islam, M. M., Y. Ishibashi, A. C. Nakagawa, Y. Tomita, M. Iwaya-Inoue, S. Arima, and S. H. Zheng. 2016, Nitrogen redistribution and its relationship with the expression of GmATG8c during seed filling in soybean: J Plant Physiol, v. 192, p. 71–74.

Marrou H, R. J. J. G. 2017, Is nitrogen accumulation in grain legumes responsive to growth or ontogeny: Physiologia Plantarum, v. 162, p. 109–122.

Mesejo, C., A. Martínez-Fuentes, C. Reig, and M. Agustí. 2019, The flower to fruit transition in Citrus is partially sustained by autonomous carbohydrate synthesis in the ovary: Plant science (Limerick), v. 285, p. 224–229.

Miyazaki, Y., Y. Maruyama, Y. Chiba, M. J. Kobayashi, B. Joseph, K. K. Shimizu, K. Mochida, T. Hiura, H. Kon, A. Satake, and J. Knops. 2014, Nitrogen as a key regulator of flowering in Fagus crenata: understanding the physiological mechanism of masting by gene expression analysis: Ecology letters, v. 17, p. 1299–1309.

Miyazaki, Y., Y. Maruyama, Y. Chiba, M. J. Kobayashi, B. Joseph, K. K. Shimizu, K. Mochida, T. Hiura, H. Kon, and A. Satake. 2014, Nitrogen as a key regulator of flowering in Fagus crenata: understanding the physiological mechanism of masting by gene expression analysis: Ecology Letters, v. 17, p. 1299–309.

Munier-Jolain N, L. A. S. C. 2008, Determinism of carbon and nitrogen reserve accumulation in legume seeds.: Comptes rendus - Biologies, v. 10, p. 780–787.

Seki, H., S. Sawai, K. Ohyama, M. Mizutani, T. Ohnishi, H. Sudo, E. O. Fukushima, T. Akashi, T. Aoki, K. Saito, and T. Muranaka. 2011, Triterpene Functional Genomics in Licorice for Identification of CYP72A154 Involved in the Biosynthesis of Glycyrrhizin: The Plant Cell, v. 23, p. 4112–4123.

Tu, B., C. Liu, B. Tian, Q. Zhang, X. Liu, and S. J. Herbert. 2017, Reduced abscisic acid content is responsible for enhanced sucrose accumulation by potassium nutrition in vegetable soybean seeds: J Plant Res, v. 130, p. 551–558.

Vaya, J., P. A. Belinky, and M. Aviram. 1997, Antioxidant Constituents from Licorice Roots: Isolation, Structure Elucidation and Antioxidative Capacity Toward LDL Oxidation: Free radical biology & medicine, v. 23, p. 302–313.

W, G. V. O. L. 1967, Ammonium Nutrition and Flowering of Apple Trees: Australian journal of biological ences, v. 4, p. 761–768.

Yan, B., J. Hou, J. Cui, C. He, W. Li, X. Chen, M. Li, and W. Wang. 2019, The Effects of Endogenous Hormones on the Flowering and Fruiting of Glycyrrhiza uralensis: Plants (Basel), v. 8.

You, C., L. Chen, H. He, L. Wu, S. Wang, Y. Ding, and C. Ma. 2017, iTRAQ-based proteome profile analysis of superior and inferior Spikelets at early grain filling stage in japonica Rice: BMC Plant Biology, v. 17.

Yu J, H. J. W. R. 2016, Down-regulation of nitrogencarbon metabolism coupled with coordinative hormone modulation contributes to developmental inhibition of the maize ear under nitrogen limitation: Planta, v. 1, p. 111–124.

Yu, X., X. Chen, L. Wang, Y. Yang, X. Zhu, S. Shao, W. Cui, and F. Xiong. 2017, Novel insights into the effect of nitrogen on storage protein biosynthesis and protein body development in wheat caryopsis: Journal of Experimental Botany, v. 68, p. 2259–2274.

Zhang, G., X. Wang, B. Wang, Y. Tian, M. Li, Y. Nie, Q. Peng, and Z. Wang. 2013, Fine mapping a major QTL for kernel number per row under different phosphorus regimes in maize (Zea mays L.): Theoretical and applied genetics, v. 126, p. 1545–1553.

Zhang, W., Z. Cao, Q. Zhou, J. Chen, G. Xu, J. Gu, L. Liu, Z. Wang, J. Yang, H. Zhang, and B. T. Ayele. 2016, Grain Filling Characteristics and Their Relations with Endogenous Hormones in Large- and Small-Grain Mutants of Rice: PloS one, v. 11, p. e0165321–e0165321.

Ai X M, He R Y, Xu Y Y, et al. 2017, Physiological and biochemical differences between flowering and non flowering potato varieties. Acta Agriculturae Universitatis Jiangxiensis, v. 39, p. 230–236.

Lin X J, Ma F S, Chen P, et al. 2019, Dynamics of carbohydrate content in leaves of coffee during flowering. Tropical Agricultural Sciences, v. 39 p. 5–9.

Li G Q, Zhang X Z, Zheng G Q, et al. 2012, Monitoring of C/N ratio of summer maize leaves based on canopy reflectance spectrum. Corn Science, v. 20 p. 81–85.

Jiang T H, Dan P P, Huang Z F, et al. 2016, Effects of Nitrogen Application on nitrogen absorption and accumulation in leaves and grain quality of oil Peony. Journal of Applied Ecology, v. 27 p. 3257–3263.

Luo S, Zhong Q P, Ge X N, et al. 2019, Effects of different N, P and K fertilization ratios on flower bud differentiation of Camellia oleifera. Forestry Science Research, v. 32 p. 131–138.

Mi H.L, Xu X, Li S.H, He J, et al. 2005, Dynamic change of the contents 、 distributions and ratioes of carbohydrate and total nitrogen in Cynanchum komarovii and Glycyrrhiza uralensis during the different periods of growth. Agricultural research in arid area, v. 1, p. 129–133.

Ni Y L. 2013, Study on the physiological basis of the difference of floret development in wheat and the regulation of cultivation measures. Shandong Agricultural University. p. 110

Peng F L. 2016, Study on seed setting habit and key technology of seed production of Atractylodes macrocephala. Guizhou University, p. 72.

Sha P. 2006, Relationship between endogenous hormone, water, nitrogen, phosphorus and potassium contents and seed setting of Ginkgo biloba. Guangxi University.

Sun Q C, Yang Y J, Chen N, et al. 2010, Study on morphological characteristics and C/N ratio during flower bud differentiation of radish. Northern Horticulture, v.17, p. 47–49.

Wang, J.Y.; Wang, W.Q.; Liu Y. 2003, Research progress in the biological characteristic and resource cultivation of Glycyrrhiza uralensis Fisch. World For. Res. v. 16, p. 28–32.

Wang J L, Yang M Y, Yang X Z, et al. 2012, Effects of phosphorus nutrition on growth and flowering of Chrysanthemum morifolium. Jiangsu Agricultural Sciences, v. 40, p. 156–159.

Wang P F. 2016, Effects of different fertilizer rates on Yield Related Traits of sunflower and economic benefit analysis. Inner Mongolia Agricultural University p. 67.

Wang, Q.E.; Ren, H.; Cao, X.L. 2011, Research and Utilization Statue of Licorice. Chin. Agric. Sci. Bull. v. 27, p. 290–295.

Wang W N, Lu J W, He Y Q, et al. 2011, Effects of N, P and K Fertilizers on yield, quality and nutrient uptake and utilization of rice. Rice science in China, v. 25, p. 645–653.

Wang X D, Yu Z W, Wang D, et al. 2003, Effects of potassium on carbohydrate content and starch accumulation in wheat stem and leaf sheath. Journal of Plant Nutrition and Fertilizer, v. 01 p. 57–62.

Wei M, Zhu Y L, Huo Y N, et al. 2019, Effects of potassium application on growth and flowering of Zinnia. China Agricultural Science and Technology Guide, v. 21 p. 148–153.

Wu X L, Li C S, Tang Y L, et al. 2017, Effects of Nitrogen Application on wheat yield, nitrogen use efficiency and light use efficiency. Journal of Applied Ecology, v. 28 p. 1889–1898.

Xu Z G, Huang D Y, Guo Q R, et al. 2017, Distribution pattern and dynamic change of nutrient elements in Dendrocalamus latiflorus before and after flowering. Guangxi Forestry Science, v. 46 p. 243–247.

Xu W D, Zheng C L, Wu X Z, et al. 2009, Dynamic changes of carbon and nitrogen contents in leaves of Myrica rubra during flower bud physiological differentiation. Fujian hot work technology, v. 34, p. 18–20+6.

Zhang C Q, Lian H, Ma G N, et al. 2018, Effects of phosphorus on sugar content, yield and quality of muskmelon leaves at fruiting stage. Modern Agriculture, v. 06 p. 17–19.

Zhang H N, Su Z X, Chen H B, et al. 2016, Effects of flower bud differentiation on Photosynthesis and metabolism of carbon and nitrogen in source leaves of Litchi chinensis. Guangdong Agricultural Sciences, v. 43 p. 50–55.

Zhao F X, Jiang Y M, Peng F T, et al. 2008, Characteristics of urea 15N absorption, allocation, and utilization by sweet-cherry. Journal of Applied Ecology, v. 03 p. 686–690.

Zhao J, Lv X L, Nie X, et al. 2016, Changes of carbon and nitrogen metabolism in leaves of Summer Black Grape during the second flower formation of winter bud. Heilongjiang Agricultural Sciences, v. 12 p. 82–86.

Zhou Y Y, Xi J L, Li J, et al. 2011, Effects of formula fertilization on Rice Yield and nitrogen utilization. Jiangsu Agricultural Sciences, v. 01 p. 75–78.

